# Infection and herbicide exposure implicate c-Abl kinase in α-Synuclein Ser129 phosphorylation

**DOI:** 10.1101/2025.01.29.635561

**Authors:** Marzieh Ehsani, Zeyang Sun, Alvaro Quevedo-Olmos, Gesa Rösler, Mahdi Rasa, David Holthaus, Saskia F. Erttmann, Thomas F. Meyer

## Abstract

**Background:** Parkinson’s disease is a complex multifactorial neurodegenerative disorder characterized by α-Synuclein aggregation in Lewy bodies, with phosphorylation at serine 129 (pSer129) being a critical pathological hallmark. However, the exact mechanisms by which environmental triggers lead to this disease phenotype remain poorly understood. In this study we compare the effects of an exemplary infection and a certain pesticide exposure on the generation of pSer129 α-Synuclein, with a focus on the involvement of cellular kinases in this process.

**Methods:** Two distinct environmental stressors were applied to neuronal cells: the pesticide rotenone and the well-studied gastric bacterium *Helicobacter pylori (H. pylori)*. Phosphorylation of Ser129 α-Synuclein was assessed by immunofluorescent staining and Western blotting. Cells were treated with mechanistically distinct c-Abl inhibitors, and pSer129 α-Synuclein was detected using Western blotting and activities of the upstream serine-threonine kinase were predicted by kinase profiling and Western blotting, analyzed by one-way ANOVA followed by Tukey’s multiple comparisons test. Moreover, transcriptome analyses of treated cells were performed and ingenuity pathway analysis and Deseq2 were applied to unravel the affected neurodegenerative pathways.

**Results:** The functional analysis of our RNA sequence data demonstrated that both *H. pylori* and rotenone induced oxidative stress and neuroinflammation by stimulating neurodegenerative pathways. Rotenone and *H. pylori* activated c-Abl, likely through the induced oxidative stress and promoted α-Synuclein phosphorylation. The kinase inhibitors Ponatinib and Asciminib effectively prevented pSer129 α-Synuclein accumulation and reversed associated gene expression changes induced by rotenone or *H. pylori*. Moreover, GSK3β appeared to be involved in the induction of Ser129 phosphorylation via activated c-Abl. Furthermore, *H. pylori’s* vacuolating cytotoxin appeared to play a crucial role in the phosphorylation of pSer129 α-Synuclein by c-Abl.

**Conclusions:** These findings highlight the pivotal role of c-Abl in α-Synucleopathies and provide insights into shared mechanisms between infection and pesticide exposure, offering potential therapeutic targets for Parkinson’s disease and related pathologies involving α-Synuclein modification.

## Introduction

Parkinson’s disease (PD) is a progressive neurodegenerative disorder characterized by the accumulation of α-Synuclein (α-Syn) aggregates within Lewy bodies in the brains of affected individuals [1]. Phosphorylation at serine 129 (pSer129) appears to be the predominant post-translational modification of α-Syn, promoting formation of α-Syn aggregates and representing a critical hallmark of PD pathology [1, 2]. This modification not only facilitates α-Syn aggregation in PD but is also implicated in other α-Synucleinopathies, including multiple system atrophy and amyotrophic lateral sclerosis [3, 4]. Notably, elevated levels of pSer129 α-Syn have been observed in cerebrospinal fluid and postmortem brain samples, suggesting its utility as a biomarker for disease progression and therapeutic monitoring [5].

PD can result from genetic and environmental factors, including exposure to toxic pesticides. Notably, rotenone has been implicated in the so-called “PD pandemic” over the past two decades [6]. As a naturally occurring pesticide, rotenone crosses the blood-brain barrier and exerts neurotoxic effects by inhibiting mitochondrial complex I, resulting in mitochondrial dysfunction, reduced ATP production and increased oxidative stress [7, 8]. These effects culminate in the death of dopaminergic neuron, a defining feature of PD [8]. Rotenone is also known to induce pSer129 α-Syn phosphorylation, which further promotes aggregation and neurodegeneration [9]. However, the mechanisms underlying this phosphorylation remain incompletely understood.

One of the key kinases that has emerged as a key player in progression of neurodegenerative diseases is c-Abl (short for cellular Abelson tyrosine kinase), a stress-activated non-receptor tyrosine kinase [10]. Under oxidative stress, c-Abl undergoes autophosphorylation at tyrosine residues Y245 and Y412, triggering its own activation [11, 12]. Once activated, c-Abl modulates several cellular pathways, including those involved in mitochondrial function that are dysregulated in PD [13]. c-Abl directly phosphorylates key proteins involved in PD pathology, including α-Syn [10, 14] and parkin [15], thus exacerbating neuronal damage. The development of c-Abl inhibitors, such as Ponatinib, has opened promising therapeutic avenues for PD, with initial studies demonstrating their potential to slow disease progression and alleviate symptoms [16, 17].

Emerging evidence underscores the role of gut microbes and dysbiosis in the development of PD [18]. Among these, the gastric pathogen *Helicobacter pylori* (*H. pylori*), which colonizes the stomach in approximately 50% of the global population, serves as an illustrative model for studying the connection between gastrointestinal inflammation and neuroinflammation [19–23]. The innervation of the gastric mucosa by the vagus nerve provides a plausible pathway for gut-brain axis signaling, linking peripheral infections to effects in the central nervous system [24–26]. Anecdotal observations and clinical studies have linked *H. pylori* infections, particularly in cases of chronic gastritis and gastric ulcers, to an increased risk of PD [19, 20]. Mechanistically, *H. pylori* induces significant inflammation by secreting pro-inflammatory molecules and generation of oxidative stress [27, 28]. Chronic inflammation can establish conditions that promote the activation of c-Abl, a key feature of the pathogen’s effect on host cell signaling [29–31]. However, the mechanisms underlying potential neuroinflammation by *H. pylori* remain poorly understood. Several virulence factors, such as the type IV secretion system [32], and effector protein cytotoxin-associated gene A (CagA), and the vacuolating cytotoxin VacA, are known to induce oxidative stress and are likely candidates for c-Abl activation and its downstream effects.

In this paper, we aimed to investigate the mechanisms by which rotenone and *H. pylori* induce α-Syn phosphorylation at Ser129 and observed that both stimuli point to a common central role of the tyrosine kinase c-Abl in this process. Using human neuroblastoma cells, we further identified the involvement of Glycogen Synthase Kinase-3 (GSK3β), a serine-threonine kinase downstream of c-Abl, likely being responsible for the direct phosphorylation of Ser129. By uncovering this convergent pathway linking environmental toxins and microbial infections to α-Synucleinopathies, this work offers insights into novel, generalized therapeutic strategies for PD.

## Results

### *H. pylori* and rotenone induce pSer129 α**-Syn**

In mouse models of PD, rotenone has been shown to induce phosphorylation of serine 129 (pSer129) α-Syn [33]. Here, we used SH-SY5Y, a human neuroblastoma cell line, which expresses several markers of peripheral neuronal progenitors and is often used as a model for neurodegenerative disease to investigate modifications of α-Syn. We treated the cells with 10 µM rotenone or infected with *H. pylori* at multiplicity of infection (MOI) 100. Upon infection, phosphorylation of CagA at Y99 occurs intracellularly in the host cell, indicating a successful infection (Additional file 1). Surprisingly, infection with *H. pylori* induced pSer129 α-Syn significantly even after 3 hours (Fig. 1A). However, phosphorylation of Ser129 culminated 6 h after rotenone treatment (Fig. 1B). High resolution confocal microscopy confirmed pSer129 α-Syn in response to both *H. pylori* and rotenone (Fig. 1C). Thus, similar to rotenone, *H. pylori* infection promotes the induction of pSer129 α-Syn in SH-SY5Y cells.

**Figure 1.**
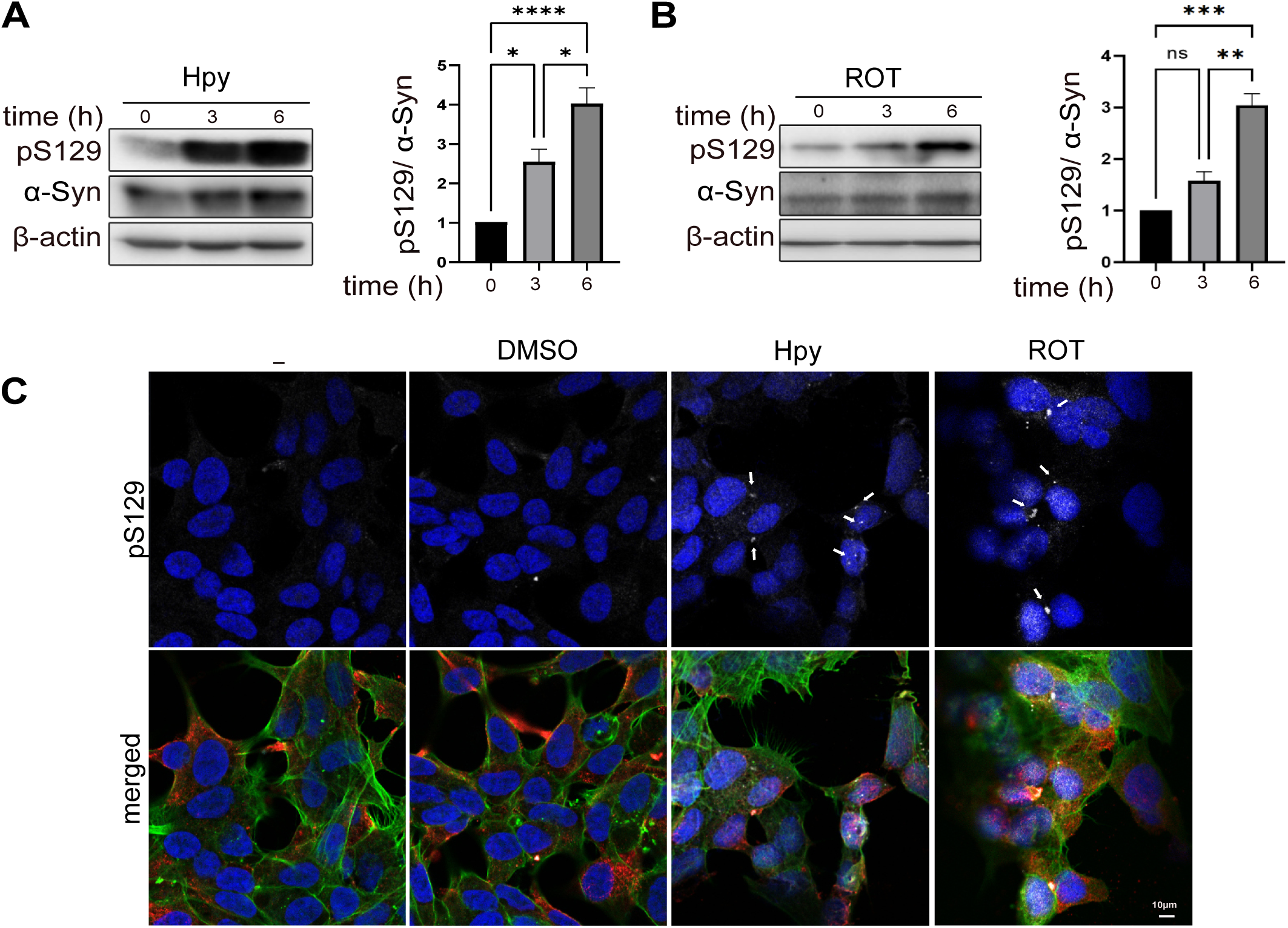
*H. pylori* and rotenone induce pSer129. α**-Syn in human SH-SY5Y.** (A) SH-SY5Y cells were infected with *H. pylori* (Hpy) at MOI 100 and total protein was extracted after 6 h and pSer129 α-Syn is detected. Quantification of pSer129 α-Syn protein bands normalized to total α-Syn is shown. (B) SH-SY5Y cells were treated either with 10 µM rotenone (ROT) or 0.1% DMSO up to 6 h and Western blotting has been performed on extracted total protein. Quantification of pSer129 α-Syn normalized to total α-Syn (C) SH-SY5Y cells were infected with *H. pylori* at MOI 100 or treated with 0.1% DMSO or 10 µM ROT for 6 h. Cells were fixed with 4% PFA and stained for total α-Syn (syn211, red), pSer129 (white), phalloidin (green) and nucleus (blue). The scale bar indicates 10 µm. The data represent n=3 independent biological replicates. Data are presented as means ± standard error of the mean (SEM). Statistical analysis was performed using one-way ANOVA followed by Tukey’s multiple comparisons test. ns, not significant; * p[<[0.05; ** p[<[0.01; *** p[<[0.001, **** p[<[0.0001.

### *H. pylori* and rotenone induce oxidative stress, inflammation and activation of c-Abl

To investigate the common pathways that are triggered by both *H. pylori* and rotenone in SH-SY5Y cells, bulk RNA sequencing was performed (GEO Accession number GSE87385). Gene set enrichment analysis (GSEA) revealed common functionally relevant pathways induced by rotenone and *H. pylori*, including stress responses, the activation of p38 mitogen-activated protein kinases (p38 MAPK), nuclear factor kappa B (Nf-κB), activator protein-1 (AP-1), Jun N-terminal kinase (JNK), activating transcription factor-2 (ATF2) and the production of inflammatory cytokines such as tumor necrosis factor alpha (TNFα), interleukin (IL)-6, IL-8, and IL-2 (Fig. 2A, Additional file 2). The cumulative impacts of increased inflammation and prolonged stress apparently led to cell death, as evident among the significantly induced pathways (Fig.2A).

**Figure 2.**
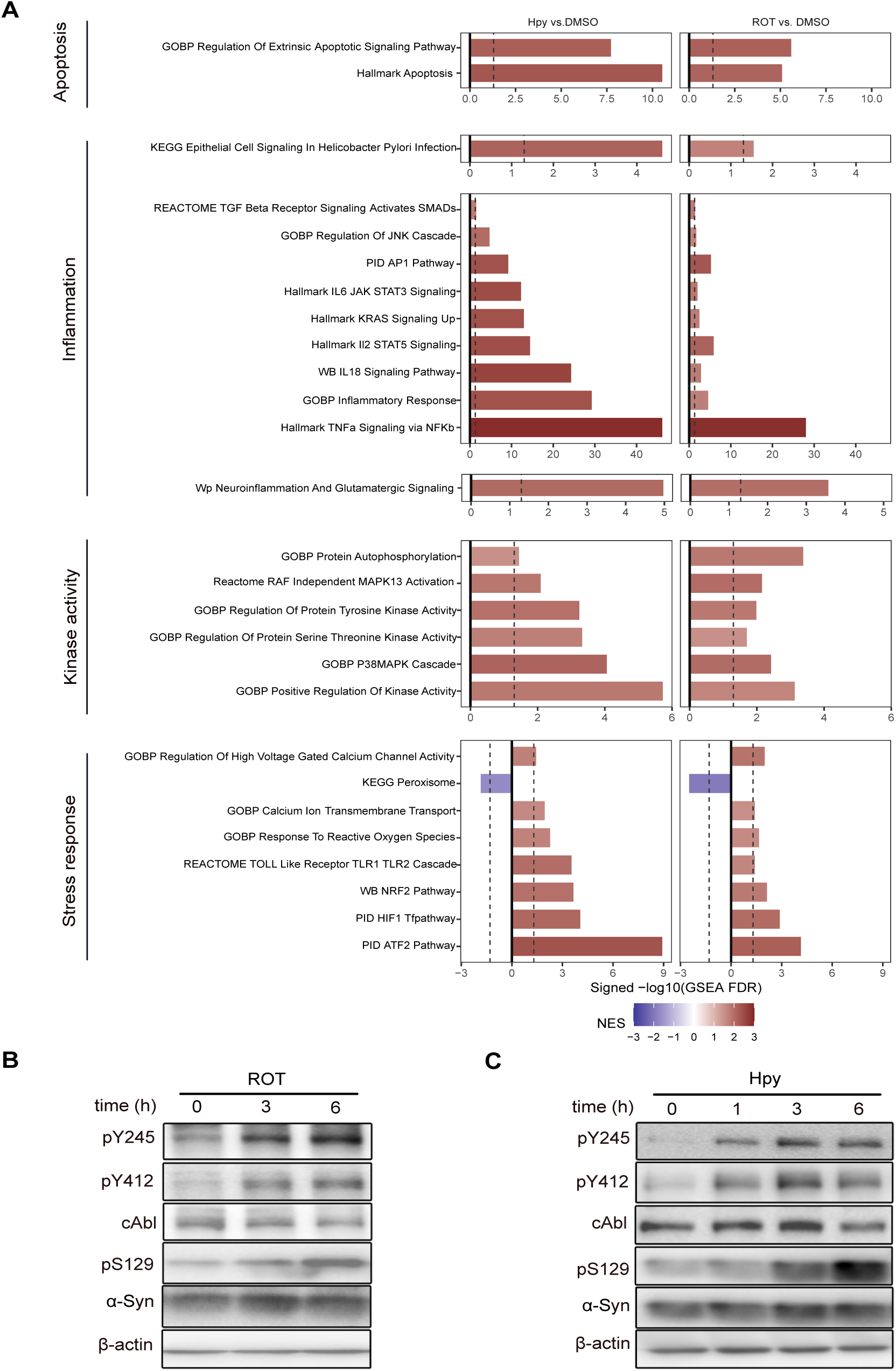
*H. pylori* and rotenone induce oxidative stress, inflammatory response and activation of tyrosine kinases. (A) SH-SY5Y cells were treated with 10 µM rotenone (ROT) or infected with *H. pylori* (Hpy) at MOI 100 for 4 h and bulk RNA sequencing is performed on the total extracted RNA. The functionally relevant significant pathways are shown for *H. pylori* and ROT vs. the control samples. The length of the bars is proportional to the FDR adjusted GSEA significances. Colors represent the Normalized Enrichment Score (NES): Red represents pathway activation and blue represent pathway inhibition compared to control samples. (B) SH-SY5Y cells were treated either with 10 µM ROT or 0.1% DMSO up to 3 and 6 h and Western blotting was performed on extracted total protein. (C) SH-SY5Y cells were infected with *H. pylori* at MOI 100 and Western blotting was performed on extracted total protein after 1, 3 and 6 h incubation.

As it is shown by the RNA sequencing results, the hallmark response of rotenone and *H. pylori* is the induced intracellular stress, which is sufficient to activate tyrosine kinase c-Abl [34]. c-Abl is a key tyrosine kinase that mediates oxidative stress in host cells upon response to infection with *H. pylori* [35, 36]. Considering the emerging role of c-Abl in initiation and progression of PD, we aimed to investigate c-Abl activity upon infection with *H. pylori* or treatment with rotenone. Enhanced phosphorylation of the two main autophosphorylation sites of c-Abl (Y412 and Y245) suggested increased kinase by both triggers (Fig. 2B and C), corresponding to the accumulation of pSer129 α-Syn in a time-dependent manner. Thus, both triggers, *H. pylori* and rotenone, induce c-Abl autophosphorylation and pSer129 α-Syn accumulation.

### *H. pylori* and rotenone induce pSer129 **α**-Syn through c-Abl

Our findings demonstrated that both *H. pylori* infection and rotenone treatment activate c-Abl and induce pSer129 α-Syn. To determine whether Ser129 phosphorylation of α-Syn is dependent on c-Abl, cells were pre-treated with Ponatinib, a third-generation c-Abl inhibitor, prior to exposure to rotenone or *H. pylori*. Infected cells exhibited strong induction of pSer129, pY412 and pY245 compared to control cells, and addition of Ponatinib effectively rescued pSer129 and c-Abl autophosphorylations (Fig. 3A). Similarly, rotenone-treated cells showed significant induction of pSer129, pY412 and pY245. In contrast, combination treatment with Ponatinib rescued c-Abl autophosphorylation and Ser129 phosphorylation (Fig. 3B).

**Figure 3.**
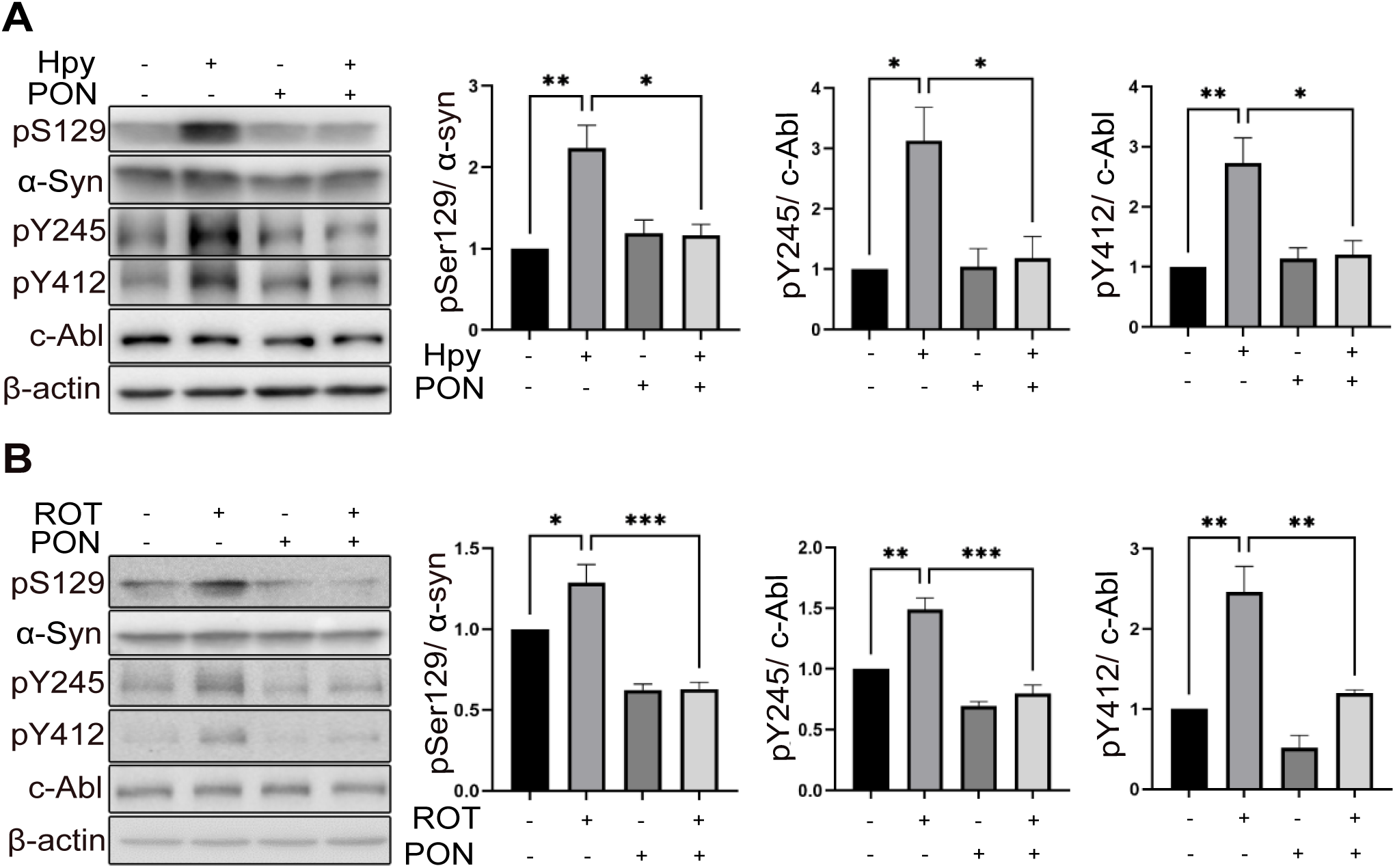
*H. pylori* and rotenone induce phosphorylation of Ser129. α**-Syn through activation of c-Abl in SH-SY5Y cells.** (A) SH-SY5Y cells were infected with *H. pylori* at MOI 100 or treated with 200 nM Ponatinib (PON) and total protein was extracted after 6 h. Quantification of pSer129 α-Syn pY245 c-Abl and pY412 c-Abl protein bands normalized to total protein is shown. (B) SH-SY5Y cells were treated either with 10 µM ROT, 50 nM PON (PON) or 0.1% DMSO up to 12 h and Western blotting was performed on the extracted total protein. Quantification of pSer129 α-Syn, pY245 c-Abl and pY412 c-Abl protein bands normalized to total protein is shown. The data represents as n=3 replicates. Means ± SEM are shown. Statistical analysis was performed using one-way ANOVA followed by Tukey’s multiple comparisons test. ns, not-significant; * p[<[0.05; ** p[<[0.01; *** p[<[0.001.

To further strengthen our data, we utilized Asciminib, a recently developed tyrosine kinase inhibitor that targets the myristate site of c-Abl [37]. Consistent with the effects of Ponatinib, Asciminib also strongly rescued Ser129 α-Syn phosphorylation and c-Abl autophosphorylations upon infection with *H. pylori* or treatment with rotenone (Additional file 3).

Thus, by employing mechanistically distinct c-Abl inhibitors, our findings demonstrated that c-Abl regulates Ser129 α-Syn phosphorylation in SH-SY5Y cells. These results confirms that *H. pylori* and rotenone induce Ser129 α-Syn phosphorylation through a mechanism dependent on c-Abl activity, highlighting the pivotal role of c-Abl kinase in this process.

### c-Abl is a major regulator of gene expression in response to *H. pylori* and rotenone

To further characterize the role of c-Abl, bulk RNA sequencing analyses were performed on SH-SY5Y cells exposed to *H. pylori* and rotenone in the presence or absence of the c-Abl inhibitor, Ponatinib. Principal component analysis (PCA) was conducted using transcripts per million (TPM) values across all genes to identify common trends in gene expression after infection with *H. pylori* or treatment with rotenone. The PCA plot shows that Ponatinib-treated samples overlap with control group (Fig. 4A), reflecting the minimal impact of Ponatinib in the absence of rotenone or *H. pylori*. Interestingly, both *H. pylori* and rotenone caused a shift in the gene expression profiles of SH-SY5Y cells (shown by arrow a), resulting in divergence of samples in similar directions along PC1 (Fig. 4A). In contrast, treatment with Ponatinib in combination with rotenone or *H. pylori* partially rescued the induced shift in gene expression (shown by arrow b), bringing them closer to the control group (Fig. 4A).

**Figure 4.**
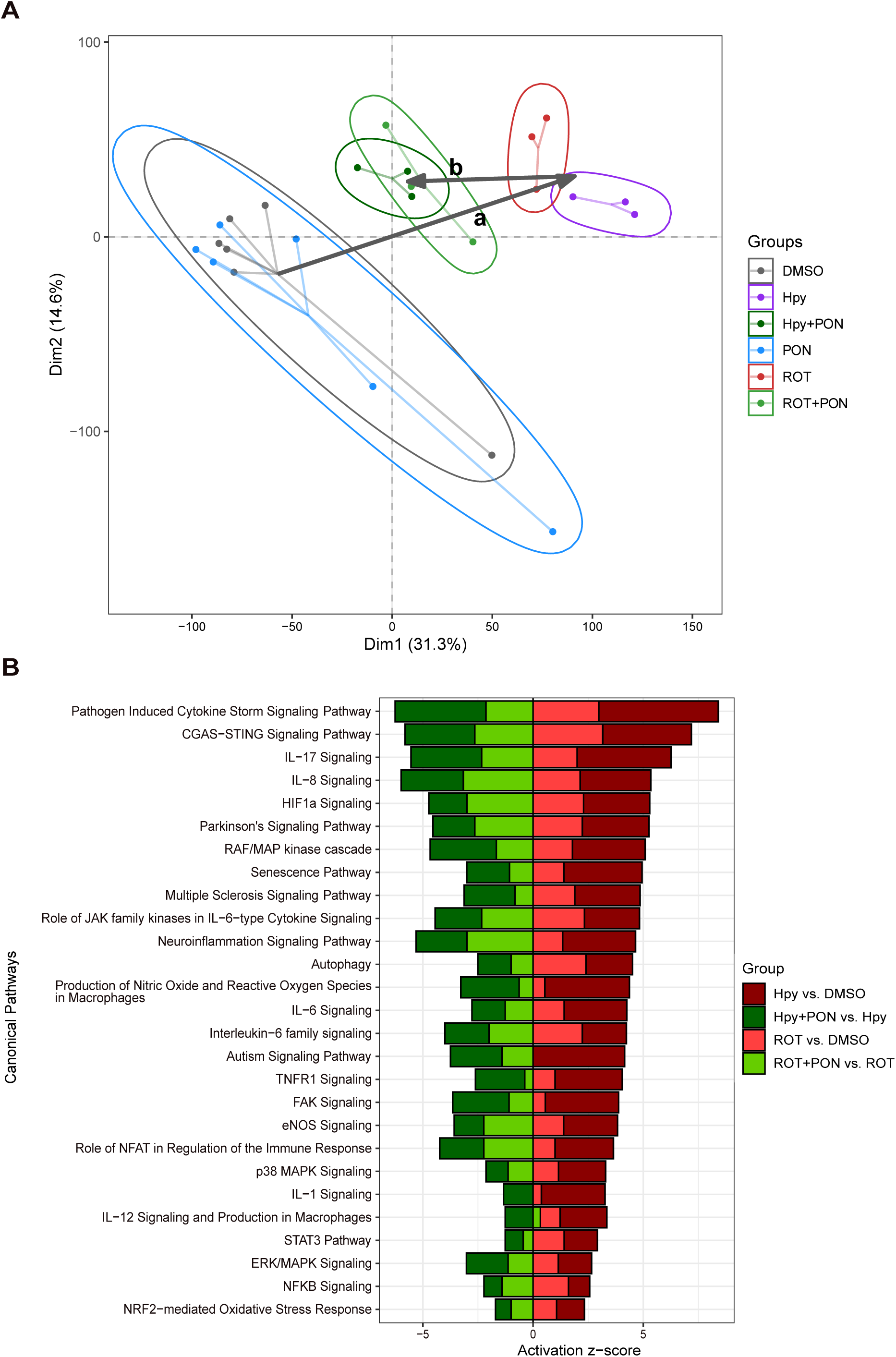
Inhibition of c-Abl rescues the induced gene expression patterns by both *H. pylori* and ROT in a similar trend. (A) SH-SY5Y cells were infected for 4 hours with *H. pylori* at MOI 100 or pre-treated one hour prior to infection with 200 nM PON or cells were treated with 10 µM ROT for 4 h or pre-treated 1 hour with 50 nM PON. For pre-treatment of and total RNA was extracted and sequenced and the gene expression patterns were shown in a PCA, plotted based on scaled TPM counts. (B) Activity prediction of the functionally selected significantly regulated canonical pathways shared by *H. pylori* and ROT, rescued by PON in the combination treatment. Colour shows the comparisons. Arrow ‘a’ indicates induction by *H. pylori* and ROT and arrow ‘b’ indicates rescue by PON. The x-axis shows the activity prediction (Z-score) for each pathway (using RNA-seq data, calculated by IPA). Positive Z-score shows activation and negative Z-score shows inhibition.

To unravel the common pathways rescued by c-Abl inhibition, ingenuity pathway analysis (IPA) was performed. The IPA results not only confirmed the induction of multiple inflammatory, oxidative stress and apoptotic pathways but also showed induction of several additional pathways upon exposure to rotenone or *H. pylori* compared to controls (Fig. 4B). Both rotenone and *H. pylori* triggered pathways associated with neuroinflammation and PD and these inductions were diminished when c-Abl was inhibited by Ponatinib (Fig. 4B).

Polo-like kinases (PLKs) have been identified as direct regulators of α-Syn phosphorylation at Ser129 and are therefore considered as potential therapeutic targets for PD [38–40]. In this study, we demonstrated that exposure to *H. pylori* and rotenone enhanced the gene expression of PLK2 and PLK3. Furthermore, *H. pylori*-induced gene expression of PLK2 and PLK3 was rescued by Ponatinib. In contrast, Ponatinib selectively rescued only PLK3 gene expression induced by rotenone (Additional file 4).

In summary, IPA suggests that c-Abl inhibition via Ponatinib has a significant rescuing effect on gene expression patterns and pathways disrupted by rotenone and *H. pylori* in SH-SY5Y cells, including those related to neuroinflammation and PD-related pathways.

### *H. pylori*- and rotenone-induced Ser129 phosphorylation involves GSK3**β** activation

To identify serine-threonine kinases (STKs) capable of phosphorylating α-Syn at Ser129, we utilized a kinome array to screen the activities of 144 STKs in rotenone-treated SH-SY5Y cells, with or without Ponatinib. The PamGene panel predicts kinase activity by measuring the phosphorylation of peptide substrates immobilized on the PamGhip microarray [41]. Upstream kinase analysis revealed that rotenone, compared to DMSO, induced the activity of a wide range of STKs including members of CMGC, Calcium/Calmodulin-dependent Kinase (CAMK), casein kinase 1 (CK1) and AGC families (Fig. 5A). Interestingly, co-treatment with Ponatinib rescued several STKs particularly MAPKs and CMGC family kinases including cyclin-dependent kinases (CDKs), MAPK11 and GSK3s (GSK3α and GSK3β) (Fig. 5B, additional file 5 and 6). In particular, dysregulation of GSK3β, a serine-threonine kinase that co-localizes with pSer129 α-Syn, has been described in PD and Alzheimer’s disease [42, 43]. To determine whether phosphorylation of Y216-GSK3β, a marker for GSK3β activation, is dependent on c-Abl activity, we analyzed its level upon rotenone stimulation and *H. pylori* infection. Our results indicate that both *H. pylori* and rotenone moderately induced pY216 GSK3β (Fig. 6A-D) and inhibition of c-Abl using Ponatinib and Asciminib reduced pY216 level (Fig. 6A-D), confirming our STKs profiling results (Fig. 5A and B).

**Figure 5.**
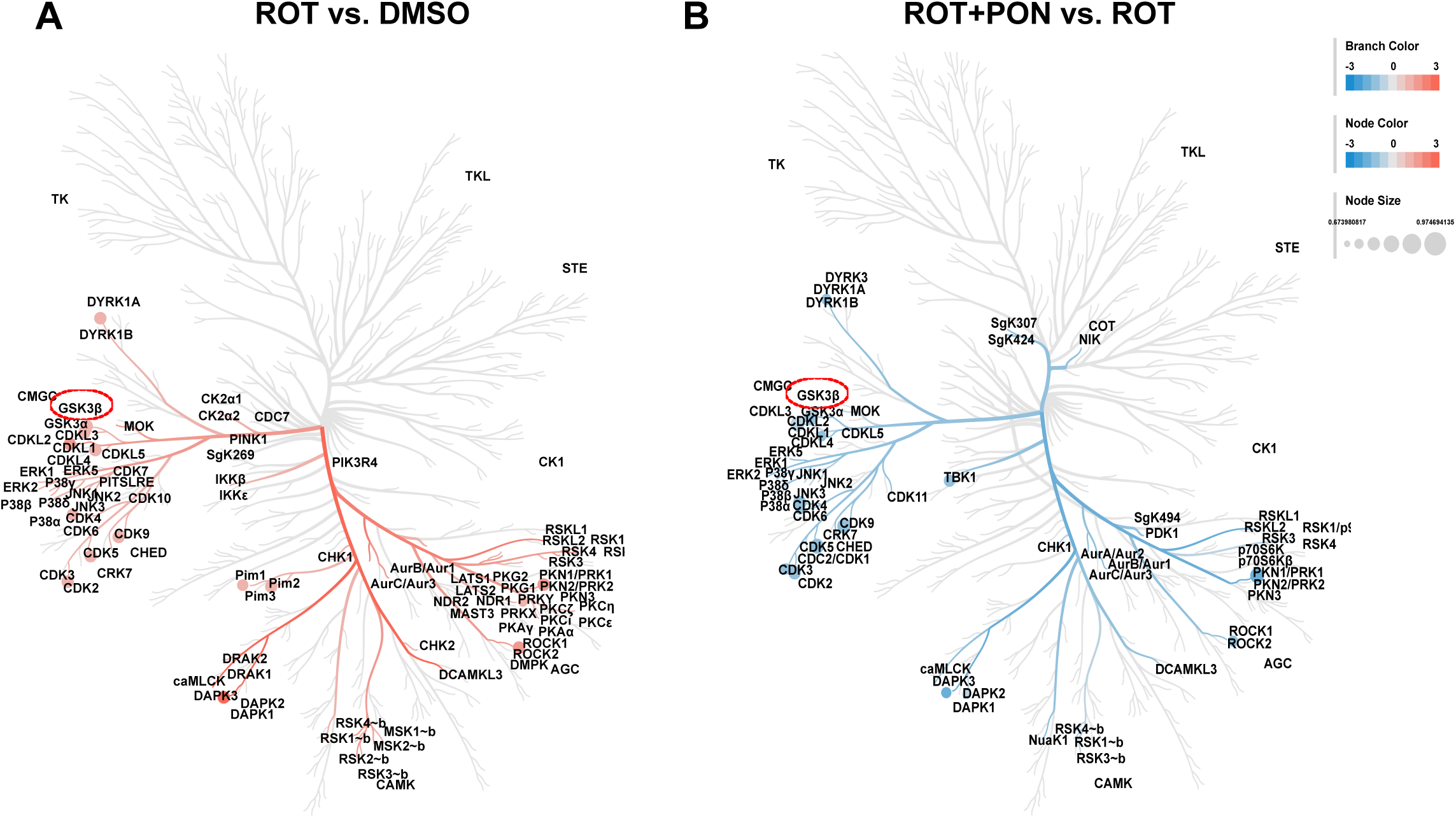
Inhibition of c-Abl inhibits several serine-threonine kinases (STKs) in rotenone-treated SH-SY5Y cells shown by STKs profiling. (A and B) SH-SY5Y cells were treated in the same way as described in Fig. 3G and total protein was extracted after 4 h and serine-threonine kinase (STKs) profiling was performed using Pamgene chip and upstream kinase analyses were conducted. The red color indicates induction of activity by rotenone (ROT) vs. DMSO and the blue color indicates reduction of activity by 50 nM Ponatinib (PON) compared to 0.1% DMSO.

**Figure 6.**
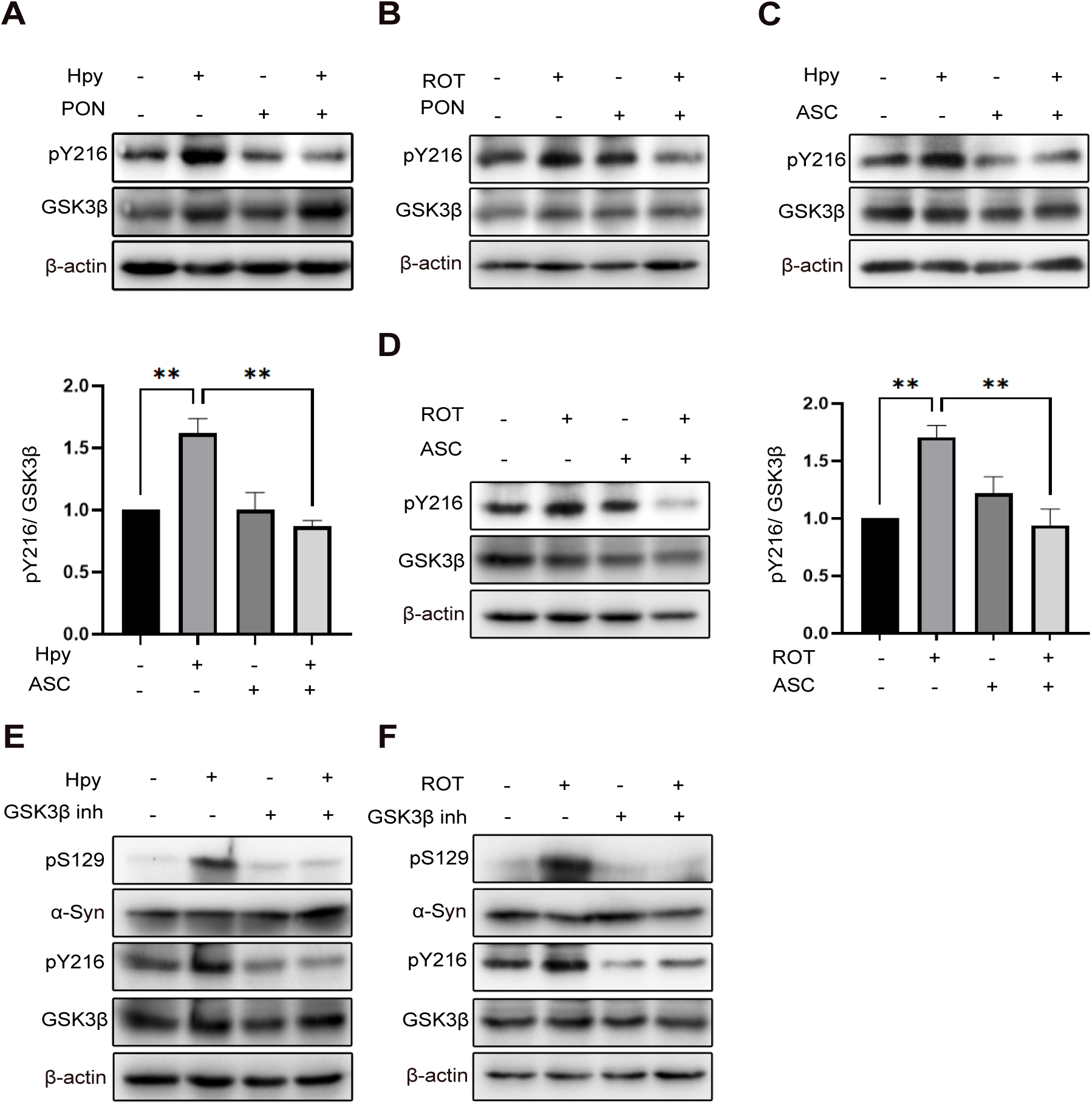
Inhibition of GSK3β inhibits pSer129 α-Syn. (A) Protein samples were prepared as described in Figure 3A and WB was performed. (B) Protein samples were prepared as described in Figure 3B and WB was performed. SH-SY5Y cells were pre-treated with 10µM Asciminib for 1 hour and infected with *H.pylori* (Hpy) WT MOI100 (C) or treated with rotenone (ROT) (D). Total protein was extracted and Western blotting was performed to detect the p-GSK3β level. The intensity of p-Y216-GSK3β was normalized to total GSK3β (E) SH-SY5Y cells were infected with *H. pylori* at the MOI 100 and pre-treated with 10 µM GSK3β inhibitor. (F) SH-SY5Y cells were treated with 10 µM ROT or Asciminib (ASC). The data represents as n=3 replicates. Means ± SEM are shown. Statistical analysis was performed using one-way ANOVA followed by Tukey’s multiple comparisons test. ns, not-significant; * p[<[0.05; ** p[<[0.01; *** p[<[0.001.

To explore the role of GSK3β in the phosphorylation of Ser129, we pre-treated the infected cells with GSK3β inhibitors and assessed pSer129 levels. Interestingly, inhibition of GSK3β in the infected cells strongly rescued the phosphorylation of Ser129 to the same level as in non-infected cells (Fig. 6E). Similarly, inhibition of GSK3β in the rotenone-treated cells also reduced the pSer129 level compared to the basal level of pY216-GSK3β in non-infected cells (Fig. 6F).

Thus, our data demonstrated that GSK3β plays a potential role in the phosphorylation of Ser129 α-Syn in *H. pylori*-infected and rotenone-treated neuroblastoma cells.

### Virulence factor VacA of *H. pylori* contributes to the phosphorylation of Ser129

*H. pylori* possesses several well-defined virulence factors that contribute to its pathogenicity [28, 44, 45]. To explore which of these virulence factors of *H. pylori* contribute to phosphorylation of Ser129 α-Syn, SH-SY5Y cells were infected with wild type (WT) *H. pylori* or mutant strains, including *H. pylori* ΔcagA (deletion of Cytotoxin-Associated Gene A), *H. pylori* ΔcagPAI (deletion of the cag Pathogenicity Island encoding both CagA and the type IV secretion system, as well as *H. pylori* ΔvacA (deletion of Vacuolating Cytotoxin A). phosphorylated α-Syn and the activity of the beforementioned kinases were tested by Western blotting. Infection with *H. pylori* WT induced pSer129 α-Syn in neuroblastoma cells, while phosphorylation of Ser129 α-Syn was reduced in response to infection with *H. pylori* ΔvacA (Fig. 7A). In contrast, deletion of cagA or the cagPAI did not significantly influence the phosphorylation level of α-Syn (Fig. 7A). To further investigate the role of VacA in the induction of pSer129 α-Syn and activation of c-Abl and GSK3β, we infected cells with *H. pylori* WT and *H. pylori* ΔvacA for different durations. Infection with *H. pylori* ΔvacA significantly reduced phosphorylation of Ser129 α-Syn, and c-Abl at both autophosphorylation sites (Y412 and Y245), and slightly affects pY216 GSK3β (Fig. 7B, C) as compared to infection with *H. pylori* WT. These findings highlight a pivotal role for VacA in the phosphorylation of Ser129 α-Syn (Fig 7D).

**Figure 7.**
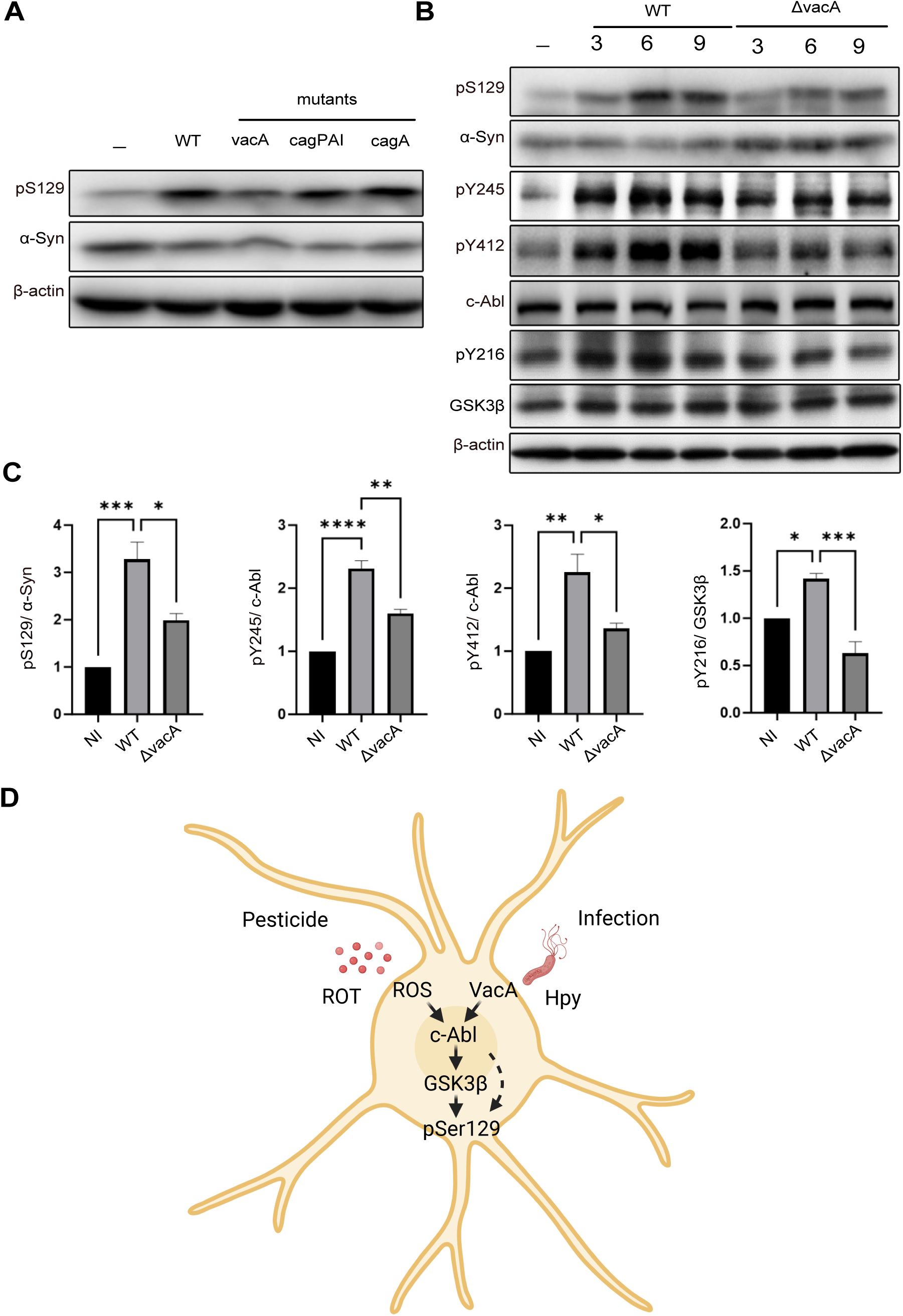
Deletion of vacA reduces c-Abl autophosphorylation and phosphorylation of GSK3β and Ser129. (A) Cells were infected for 6 hours with various strains of *H. pylori* at an MOI of 100, including P12 (WT), P12 ΔcagA (cagA mutant), P12 ΔcagPAI, and P12 ΔvacA. Total proteins were then extracted, and Western blot analyses were performed. (B) Cells were infected with *H. pylori* (MOI 100); P12 (WT) or P12 ΔvacA and total proteins were extracted at 3, 6, and 9 h post-infection. (C) Quantifications of pS129 α-Syn, pY245 c-Abl, pY412 c-Abl and pY216 GSK3β normalized to the total protein levels at time point 6 hours. (D) Schematic illustration depicting different environmental stressors leading to c-Abl activation and subsequent induction of pSer129 α-Syn. Created with Biorender.com. The data represents as n=3 replicates. Means ± SEM are shown. Statistical analysis was performed using one-way ANOVA followed by Tukey’s multiple comparisons test. ns, not-significant; * p[<[0.05; ** p[<[0.01; *** p[<[0.001.

## Discussion

Phosphorylation of serine 129 (pSer129) on α-Syn is a well-established molecular hallmark of PD. In this study, we identify the tyrosine kinase c-Abl as a central signaling hub that mediates Ser129 phosphorylation in response to diverse environmental stressors, including the pesticide rotenone and infection with the bacterial pathogen *H. pylori*. These findings shed new light on the molecular pathways connecting these distinct triggers to the modification of Ser129 on α-Syn. c-Abl activation, identified by autophosphorylation at Y245 and Y412, is triggered by oxidative stress conditions induced by both rotenone and *H. pylori* [36, 46]. This supports the idea that oxidative stress is a critical driver of c-Abl activation in neurodegenerative diseases [10, 47]. Once activated, c-Abl disrupts key cellular processes, including mitophagy and apoptosis regulation, contributing to the aggregation of α-Syn [10, 13]. Here, we demonstrated that pharmacological inhibition of c-Abl using Ponatinib or Asciminib effectively reduces pSer129 α-Syn phosphorylation and mitigates associated transcriptional alterations, underscoring c-Abl’s central role in PD pathology. Rotenone and *H. pylori* are both known to induce oxidative stress [48, 49]. While *H. pylori* is well-documented to activate c-Abl [31], we provide the first clear evidence that rotenone directly triggers c-Abl autophosphorylation, a robust and reproducible effect observed across various experimental conditions. Furthermore, RNA sequencing and PCA revealed that both stressors significantly impact neurodegenerative pathways. Kinome profiling identified downstream mediators such as GSK3β, a kinase known to play a critical role in phosphorylation of Ser129 α-Syn and aggregation [42]. These findings suggest a unifying mechanism by which diverse environmental and microbial stressors contribute to the amplification of α-Synucleinopathies.

While c-Abl’s role in phosphorylating α-Syn at Y39 and Y125 has been previously proposed [47], this study is the first to demonstrate its unexpected, likely indirect, involvement as a tyrosine kinase in the phosphorylation of Ser129 α-Syn. Elevated pSer129 α-Syn levels in cerebrospinal fluid and post-mortem brain tissue strongly correlate with Lewy body formation and PD severity [2, 47, 50], establishing it as a valuable biomarker for early diagnosis and monitoring of disease progression. By employing c-Abl inhibitors with distinct mechanisms of action, we confirm c-Abl’s critical role in this post-translational modification. Notably, c-Abl inhibition not only reduces pSer129 levels but also restores global gene expression patterns disrupted by rotenone or *H. pylori*, underscoring its broad regulatory influence.

Our STK analysis demonstrated that inhibition of c-Abl inhibits many kinases that are induced by rotenone, corroborating the role of c-Abl in activation of STKs. Among these c-Abl regulated STKs, the role of GSK3β in direct phosphorylation of α-Syn is well established. Our data suggest GSK3β to be one of the downstream mediators of S129 phosphorylation. GSK3β inhibition has emerged as a potential therapeutic approach in amyotrophic lateral sclerosis models by rescuing disease relevant phenotype, thus enhancing neuronal survival [51]. Collectively, the intricate interplay between rotenone-induced mitochondrial damage, c-Abl activation, α-Syn aggregation, and disrupted autophagy pathways underscores the complexity of PD pathogenesis. These findings suggest that therapeutic strategies targeting c-Abl inhibition, modulation of α-Syn phosphorylation, and restoration of autophagy could offer promising avenues for controlling PD and attenuating its progression.

Case-control studies have identified a link between *H. pylori* infection and the severity of neurodegenerative diseases, including PD [19]. One proposed mechanism is that outer membrane vesicles from *H. pylori* may cross the blood-brain barrier and deliver neurotoxic factors to the central nervous system [52]. In this study, we demonstrate that the VacA cytotoxin significantly contributes to c-Abl activation and phosphorylation of Ser129 α-Syn. Our findings reveal that deletion of vacA strongly reduces c-Abl autophosphorylation and α-Syn modification, whereas other virulence factors of *H. pylori*, such as CagA, play a less prominent role. These results underscore the neurotoxic effects of *H. pylori* VacA and highlight the importance of bacterial toxins and the gut-brain axis in PD pathology.

The observed similarities between rotenone and *H. pylori* as inducers of α-Syn phosphorylation highlight a broader mechanism that offers actionable therapeutic targets. The effectiveness of c-Abl inhibitors, particularly the highly specific Asciminib, in reducing pSer129 α-Syn levels and restoring normal transcriptional profiles underscores their potential to slow the progression of PD. Furthermore, targeting downstream kinases such as GSK3β, which mediate c-Abl-induced pSer129 phosphorylation, may provide complementary therapeutic strategies.

The interplay between environmental toxins and microbial infections emphasizes the multifactorial nature of α-Synucleinopathies. Our findings suggest that PD can result from diverse environmental and microbial stressors, necessitating an integrative research approach that combines toxicological and microbiological perspectives. By identifying c-Abl as a central signaling hub linking these stressors to α-Syn pathology, this study lays the foundation for novel therapeutic strategies in PD. Targeting c-Abl and its downstream pathways holds promise for mitigating disease progression and addressing the effects of various stressors in α-Synucleinopathies. Future research should focus on assessing the long-term efficacy of c-Abl inhibitors and their potential role in preventing disease onset.

## Material and Method

### Cell culture, starvation and treatment

SH-SY5Y cells (ATCC, CRL-2266, RRID: CVCL_0019) were maintained at 37°C in a 5% CO2 humidified incubator. SH-SY5Y cells were cultured in medium composed of 40% MEM (#31095029, Thermo Fisher), 40% Ham’s F-12 Nutrient Mix (#31765068, Thermo Fisher),15% FBS (#10,437,028, Thermo Fisher), 1× MEM Non-Essential Amino Acids Solution (#11140050, Thermo Fisher), 25mM HEPES, 1mM sodium pyruvate (#11360039, Thermo Fisher). To split the cells, accutase (#00-4555-56, Thermo Fisher) was used to detach the cells. For Hela cells, DMEM (#DMEM-HPSTA, Capricon), 10% heat-inactivated FBS and 25 mM HEPES (#HEP-B, Capricon) was used to prepare the growth medium. Prior to each experiment, cells were starved in medium containing no FBS for 2 hours. For Kinase profiling and RNA-sequencing of rotenone (#R8875, Sigma-Aldrich) treated cells, cells were treated with Ponatinib 50 nM (Seleckchem, #S1490) one hour before rotenone (10 µM) treatment. For Western blotting purposes, cells were also pre-treated with 10 µM Asciminib (#HY-104010, MedChemExpress) for 1 hour and 10 µM GSK3β inhibitor (#Cay13122-5, Biomol).

### Infection

*H. pylori* culture was prepared as described before [94]. Briefly, the P12 WT strain is enriched in bacto brain hear fusion (#237500, BD) with 10% heat-inactivated FBS is grown on hpGC agar (#CM0367, Oxoid) plate containing 10% heat-inactivated horse serum and 10% vitamin mix. Construction of *H. pylori* ΔvacA, ΔcagA and ΔcagPAI strains utilized in this study is described somewhere else [53, 54]. To grow the mutant strains, chloramphenicol (4 µg/ml) was added to the hpGC agar. Typically, bacteria were re-streaked after 2-3 days and they were used for infection 16-24 hours after re-streaking. The streaked plates were incubated at 37°C under microaerophilic conditions for 24 h. For infection experiments an MOI of 100 was used. Bacteria were plated after the infection to calculate the actual MOI. For infection experiments, cells were pre-treated with 200 nM Ponatinib and then infected with *H. pylori* at MOI 100. Additionally, cells were pre-treated with 10 µM Asciminib for 1 h and 10 µM GSK3β inhibitor.

### Western blotting and antibodies

The cells were harvested in 2× laemmli buffer and boiled at 95°C for 10 min. for a-syn detection, samples were loaded on 13.5% polyacrylamide gel and blotted onto a methanol-activated PVDF membrane (#11804131, ThermoFisher) and fixed with 4% paraformaldehyde (PFA) and 0.1% glutaraldehyde. For other targets, 10% polyacrylamide gels and nitrocellulose membranes (#15209804, Fischerscientific) were used. Protein transfer was performed in the presence of 20% methanol. Membranes were blocked in Roti-Block 1× (#T144.1, Roth). For α-Syn, #2642S cell signaling, pSer129 α-Syn, #23706S cell signaling, c-Abl #2862S cell signaling, pY245 c-Abl #07-787 sigma, pY412 c-Abl #C5240-1VL Merk, GSK3α/β #5676S cell signaling, p-GSK3 #05-413 Millipore, β-actin #A5441 sigma, anti-rabbit #7074S cell signaling, anti-mouse #7076S cell signaling were used.

### Immunofluorescent (IF) staining and confocal Microscopy

IFs were performed as it is described elsewhere [55]. Briefly, cells were washed with 1×PBS and fixed with 4% paraformaldehyde for 10 min. They were subsequently permeabilized for 30 min at RT using a permeabilization buffer composed of 0.1 M glycine, 0.2% Triton X-100, and 0.1% Tween-20 in tris-buffered saline (TBS). Cells were blocked with blocking buffer composed of 1% normal goat serum, 3% bovine serum albumin, 0.2% Triton X-100, and 0.1% Tween-20 in TBS for 3 h. The coverslips were incubated with a 1:100 dilution of primary anti-pSer129 α-Syn antibody (#23706S, cell signaling) and α-Syn syn211 antibody (#32-8100, Thermofischer) in a blocking buffer overnight. On the next day, they were washed 3x with a washing buffer (0.1% Triton X-100, and 0.05% Tween-20 in TBS) and incubated with Phalloidin A488 (#abA12379, Abcam), Hoechst 33342 (H1399, Invitrogen) and the secondary antibodies A647 anti-rabbit (#711-605-152, Jackson Immuno Research), 647 mouse (#715-605-15, Jackson Immuno Research,). Coverslips were washed 3x with with washing buffer, 1× with TBS 1× with deionized water before mounting on glass slides with Fluoromount-G (#15586276, Invitrogen). Images were taken using a cLSM 880microscope (Zeiss), equipped with Plan-Apochromat 20x/0.8 M27and C-Apochromat 40x/1.20 water M27 objectives and analyzed with ZEN blue software (v3.5; RRID : SCR_013672) and FIJI.

### mRNA-sequencing and analysis

Gene expression changes were examined acutely 4 hours after treatment. The total RNA was extracted using RNeasy Plus Mini Kit (Qiagen, #74136). In this study, the sequencing of the samples was performed using the PE150 sequencing strategy on NovaSeq X Plus (Illumina, San Diego) by Novogene UK (Cambridge, UK). Messenger RNA (mRNA) was purified from total RNA using poly-T oligo-attached magnetic beads. After fragmentation, the first strand cDNA was synthesized using random hexamer primers followed by the second strand cDNA synthesis. The library was ready after end repair, A-tailing, adapter ligation, size selection, amplification, and purification.

Quality of raw reads was analyzed by *fastqc*. In order to obtain the matrix of gene counts, a custom pipeline written using the python workflow manager *Snakemake* available on https://github.com/aqvd/snakemake_workflows/blob/main/Chip-Seq/Snakefile_RNAseq.py was used. Briefly, raw reads are mapped to the Human reference genome (GRCh38) using *hisat2*. Alignments with both pairs aligned in the proper orientation are used to quantify gene expression using *htseq-count.* Gene annotation from Ensembl (release 97) was used. The matrix with gene expression counts created from *htseq-count* was used as starting point for Differential Expression Analysis using the DESeq2 R package (v 1.40.2). Genes with an FDR adjusted DESeq2 P-value <0.05 and log2 Fold Change < -1 or > 1 were classified as Up- and Down-regulated respectively. Gene Set Enrichment Analysis (GSEA) was performed on the gene list ranked by log2 Fold Changes using the *fgsea* (v 1.26.0) implementation of the algorithm implemented in the clusterProfiler (v 4.8.3) R package. PCA plots were generated after scaling the TPM normalized expression matrix using the function *prcomp* and plot using the package *factoextra* (v 1.0.7).

Moreover, for functional and pathway analysis, the Ingenuity Pathway Analysis (IPA v45868156) was performed as it is previously discussed [56]. To do this, Deseq2 differential expression results (adjusted *p* value[<[0.05) were uploaded in IPA.

### Kinome activity profiling

For measuring the activity of kinases, the PamChip peptide arrays were used as described elsewhere [41]. Briefly, kinome profiling for serine/threonine kinases in a protein lysate from SH-SY5Y cells was performed by PamGene platform[41]. The preliminary analysis of the row data has been performed by upstream kinase analysis (UKA) algorithm which generated the kinome trees. The row data was filtered using the threshold cut-off for the median final score (kinase score) > 1.2. The kinase statistic represents the log fold change scaled by the noise. The median kinase statistic<0 is considered as inhibited activity (shown in blue) and median kinase statistic>0 is considered as induced activity (shown in red). The size of the nodes is proportional to the significance score.

### Statistical analysis

Statistics for differential gene analysis was detailed above in the RNA-Seq section. Figures were plotted using Prism 10 (GraphPad) or R (v4.1.0).). For multiple comparisons, we used one-way ANOVA followed by Tukey’s multiple comparisons test, recommended by Graphpad. Differences were considered significant if p < 0.05. P values ≤ 0.05 are considered statistically significant. Asterisks indicate statistical significance values as follows: * p < 0.05,** p < 0.01, *** p < 0.001, **** p < 0.0001. Data are represented as mean ± SEM from at least three independent biological replicates. The panel composition and annotations were created using Affinity Designer 2.1.1 (Serif).

### Ethical statement

The cell lines used in this study were obtained from (ATCC, CRL-2266, RRID: CVCL_0019), and their use complied with all institutional and ethical guidelines. This study did not involve human participants, animal subjects, or other ethical concerns requiring institutional approval.

## Author’s contributions

M.E. designed and performed the experiments, analyzed the data, and wrote the manuscript. Z.S. performed experiments and analyzed the results. A.Q.O. and M.R. analyzed the data and revised the manuscript. G.R. and D.H. contributed to performing experiments and revised the manuscript. S.F.E. provided advice in designing experiments and revised the manuscript. T.F.M. supervised the project and revised the manuscript.

## Availability of data and materials

All data supporting findings of this study are available within the paper and its additional files. Further inquiries can be directed to the corresponding author.

## Acknowledgements and funding

M.E. was supported by a postdoctoral fellowship from Focus Biomed Foundation, Berlin (SFB). Z.S. was supported by a doctoral fellowship from CSC, China. T.F.M. received laboratory support from the Max Planck Society by a director emeritus appointment, through his senior professorship at CAU, and from SFB. T.F.M. is a recipient of the ERC Advanced grant MADMICS, ID 885008. The graphical abstract was created with permission from BioRender.com.

