## additional file 1 for "Infection and herbicide exposure implicate c-Abl kinase in α-Synuclein Ser129 phosphorylation"

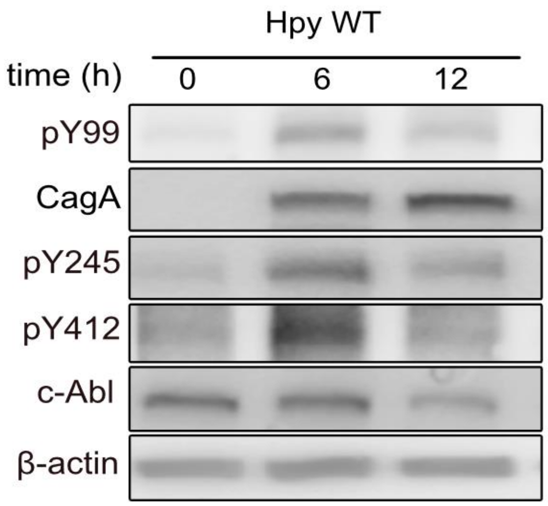


**Additional file 1.** SH-SY5Y cells were infected with *H. pylori* wild type at MOI 100. Total protein was extracted at different time points and Western blotting was performed to check phosphorylation of CagA at Y99 and autophosphorylation of c-Abl at Y412 and Y245.
