## additional file 2 for "Infection and herbicide exposure implicate c-Abl kinase in α-Synuclein Ser129 phosphorylation"

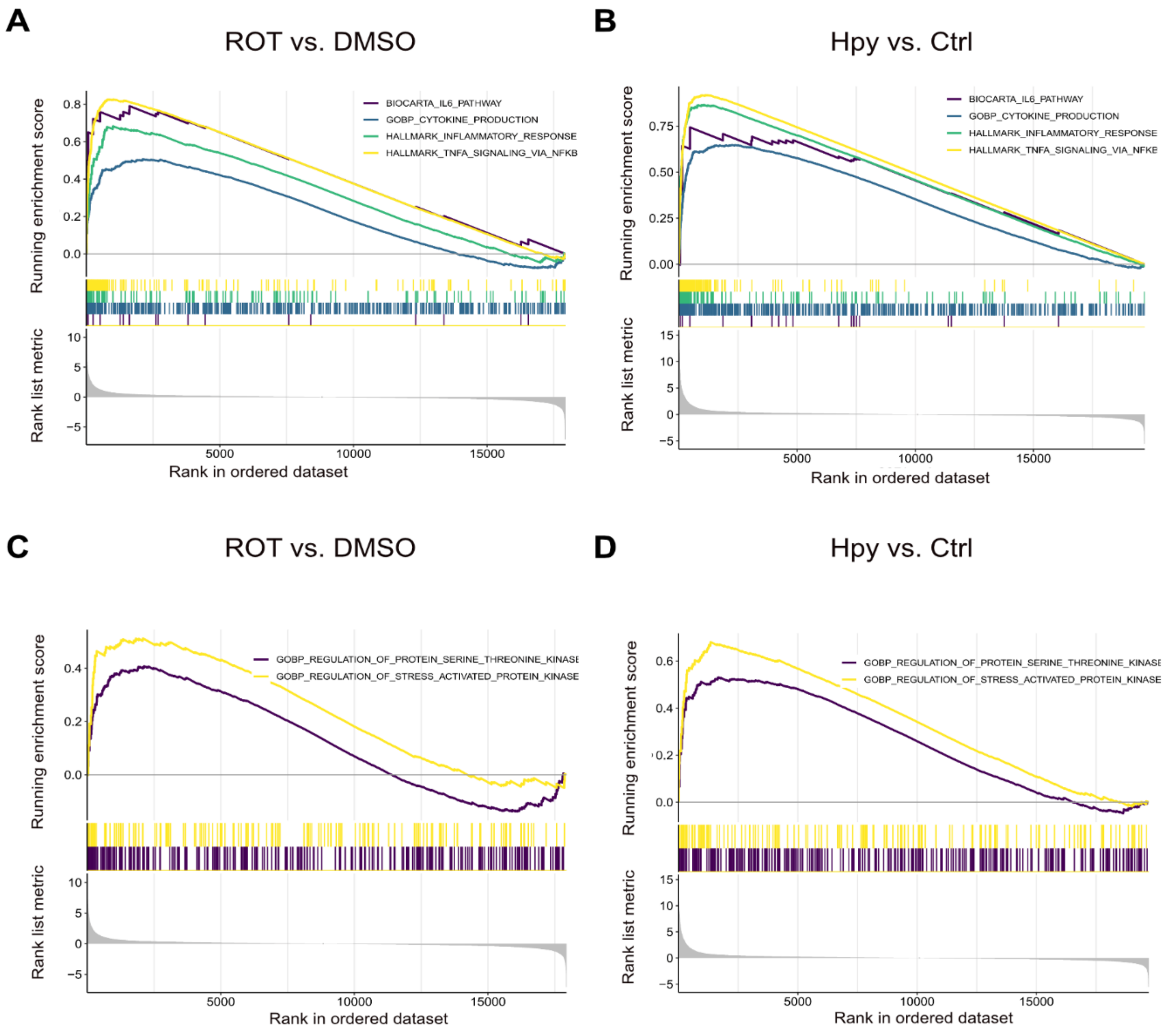


**Additional file 2.** SH-SY5Y cells were treated with rotenone (ROT) or infected with *H.pylori* (Hpy) and total RNA was extracted after 4h. Differentially expressed genes were identified using the DESeq2 R package (v 1.40.2). Gene Set Enrichment Analysis (GSEA) was performed on the gene list ranked by log2 Fold Changes using the *fgsea* (v 1.26.0) implementation of the algorithm implemented in the clusterProfiler (v 4.8.3) R package. The displayed pathways were filtered based on significance, with an adjusted p-value threshold of <0.05. A and B show enrichment of inflammatory pathways. C and D show activation of cell kinases including tyrosine and serine-threonine kinases. The data represents as n=3 replicates.
