## additional file 3 for "Infection and herbicide exposure implicate c-Abl kinase in α-Synuclein Ser129 phosphorylation"

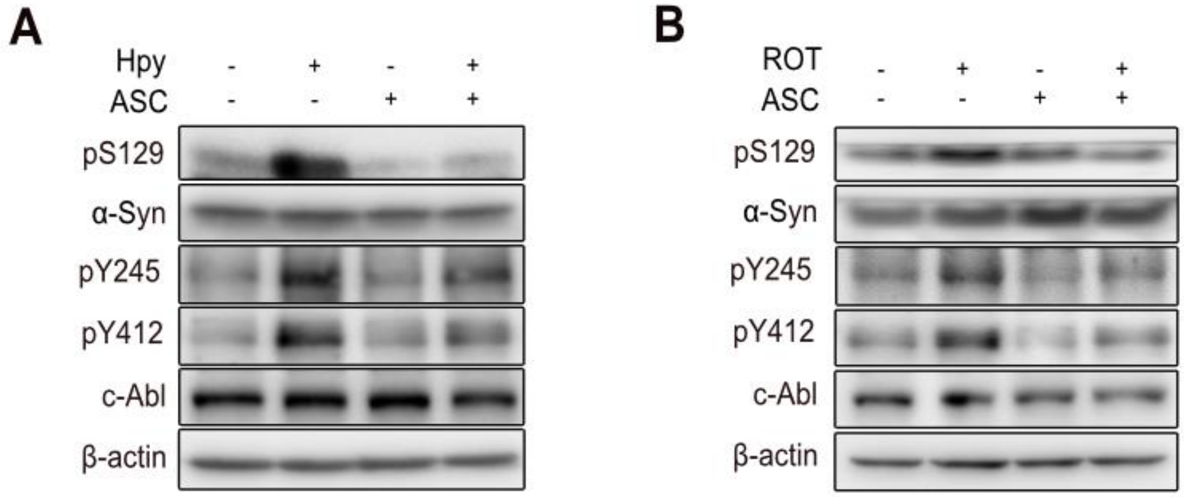


**Additional file 3.** (A) SH-SY5Y cells were infected with *H. pylori* (Hpy) at MOI 100 or pre-treated with 20 µM Asciminib (ASC) and total protein was extracted after 6h and Western blotting was performed . (B) SH-SY5Y cells were treated either with 10 µM rotenone (ROT) or 10 µM ASC and total protein was extracted after 12 h and Western blotting was performed.
