## additional file 4 for "Infection and herbicide exposure implicate c-Abl kinase in α-Synuclein Ser129 phosphorylation"

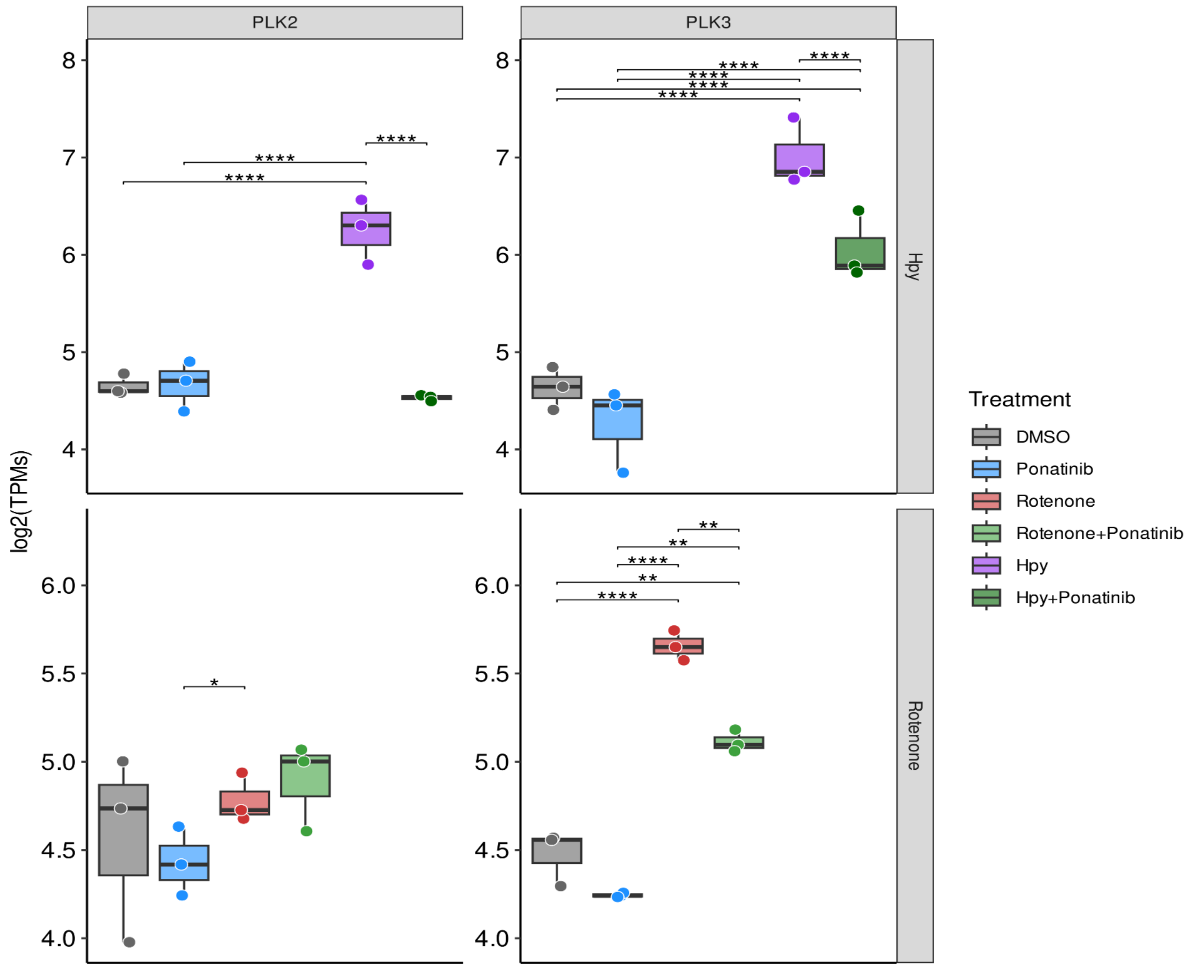


**Additional file 4.** SH-SY5Y cells were infected with *H. pylori* (Hpy) or treated with rotenone (ROT) and total RNA was extracted after 4h. Differentially expressed genes were identified and RNA counts were shown as log 2 of transcripts per million (TPM). *H. pylori* and rotenone enhanced the gene expression of Polo-like kinases2 (PLK2) and PLK3. Furthermore, *H. pylori*-induced gene expression of PLK2 and PLK3 was rescued by Ponatinib. In contrast, Ponatinib selectively rescued only PLK3 gene expression induced by rotenone. The data represents as n=3 replicates. Means ± SEM are shown. Statistical analysis was performed using Deseq2 R package. ns, not-significant; * $p < 0.05$; ** $p < 0.01$; *** $p < 0.001$.
