## additional file 5 for "Infection and herbicide exposure implicate c-Abl kinase in α-Synuclein Ser129 phosphorylation"

**Additional file 5.** The table represents upstream kinases that show induced activities upon rotenone compared to control treatment. Upstream kinase analysis (UKA) was performed on the raw data obtained from Pamgene Serine-Threonine kinase screening. The preliminary analysis has been performed by UKA algorithm. The row data to draw the kinome tree was further filtered using the threshold cut-off for the median final score (kinase score) > 1.2. The kinase statistic represents the log fold change scaled by the noise. The median kinase statistic<0 is considered as inhibited of activity and median kinase statistic>0 is considered as induced activity.

| Kinase Name | Kinase Uniprot ID | Kinase Group | Kinase Family | Max Final score |
|---|---|---|---|---|
| DAPK3 | O43293 | CAMK | DAPK | 3.673664139 |
| MYLK3 | Q32MK0 | CAMK | MLCK | 3.673664139 |
| DCLK3 | Q9C098 | CAMK | DCAMKL | 2.974694135 |
| AURKA | O14965 | CAMK | AUR | 2.71942163 |
| RPS6KC1 | Q96S38 | OTHER | RSKL | 2.673664139 |
| RPS6KA3 | P51812 | AGC-CAMK | RSK | 3.673664139 |
| PIM2 | Q9P1W9 | CAMK | PIM | 3.372634143 |
| PKN1 | Q16512 | AGC | PKN | 2.594482893 |
| MAPK7 | Q13164 | CMGC | MAPK | 2.71942163 |
| RPS6KA6 | Q9UK32 | AGC-CAMK | RSK | 3.372634143 |
| COQ8A | Q8NI60 | Atypical | ABC1 | 2.093880542 |
| RPS6KA2 | Q15349 | AGC-CAMK | RSK | 2.527536103 |
| CDKL2 | Q92772 | CMGC | CDKL | 1.910236146 |
| RPS6KA4 | O75676 | AGC-CAMK | RSK | 1.992422902 |
| PIM1 | P11309 | CAMK | PIM | 2.040195683 |
| RPS6KA1 | Q15418 | AGC-CAMK | RSK | 2.060880282 |
| CDK2 | P24941 | CMGC | CDK | 1.910236146 |
| CILK1 | Q9UPZ9 | CMGC | RCK | 1.804432419 |
| ROCK1 | Q13464 | AGC | DMPK | 1.895512889 |
| AURKB | Q96GD4 | CAMK | AUR | 1.957660795 |
| DYRK1A | Q13627 | CMGC | DYRK | 1.744245213 |
| RPS6KB1 | P23443 | AGC | RSK | 2.050414849 |
| MAPK12 | P53778 | CMGC | MAPK | 1.974694135 |
| MAPK3 | P27361 | CMGC | MAPK | 2.117361638 |
| CDKL5 | O76039 | CMGC | CDKL | 2.040195683 |
| MAPK11 | Q15759 | CMGC | MAPK | 2.030211463 |
| RPS6KA5 | O75582 | AGC-CAMK | RSK | 2.051195954 |
| CSNK2A1 | P68400 | CMGC | CK2 | 1.62834116 |
| CDK5 | Q00535 | CMGC | CDK | 1.563074429 |
| PRKAB1 | Q9Y478 | NA | NA | 1.509311283 |
| PDPK1 | O15530 | AGC | PKB | 1.480539541 |
| NUAK1 | O60285 | CAMK | CAMKL | 1.456180195 |
| MAPK1 | P28482 | CMGC | MAPK | 1.483332441 |
| MAPK9 | P45984 | CMGC | MAPK | 1.477764487 |
| MAPK10 | P53779 | CMGC | MAPK | 1.418391634 |
| MAPK8 | P45983 | CMGC | MAPK | 1.448354857 |
| GSK3A | P49840 | CMGC | GSK | 1.477764487 |
| MAPK13 | O15264 | CMGC | MAPK | 1.71942163 |
| CDK11A | Q9UQ88 | CMGC | NA | 1.385862409 |
| CDK9 | P50750 | CMGC | CDK | 1.327311165 |
| PRKY | O43930 | AGC | PKA | 1.759850287 |
| PRKCZ | Q05513 | AGC | PKC | 1.816331643 |
| CDK13 | Q14004 | CMGC | CDK | 1.413592751 |
| CDK4 | P11802 | CMGC | CDK | 1.297087182 |
| PRKCH | P24723 | AGC | PKC | 1.616759288 |
| GSK3B | P49841 | CMGC | GSK | 1.288057865 |
| MYLK2 | Q9H1R3 | CAMK | MLCK | 1.223356919 |
| CHEK1 | O14757 | CAMK | CAMKL | 1.65247484 |
| CHEK2 | O96017 | CAMK | RAD53 | 1.30019695 |
| IKBKE | Q14164 | OTHER | IKK | 1.556392843 |

| Mean Kinase Statistic | Mean Significance Score | Mean Specificity Score | Mean peptide set size |
|---|---|---|---|
| 2.949047483 | 0.974694135 | 2.698970004 | 3 |
| 2.995521184 | 0.974694135 | 2.598626672 | 4 |
| 2.624403059 | 0.974694135 | 1.94367268 | 3 |
| 1.965797985 | 0.974694135 | 1.55268161 | 5 |
| 2.415259119 | 0.974694135 | 1.614028508 | 3 |
| 1.435955963 | 0.974694135 | 1.737602422 | 21 |
| 1.238702052 | 0.974694135 | 1.592343954 | 37 |
| 1.787098431 | 0.974694135 | 1.435131873 | 7 |
| 1.403928342 | 0.974694135 | 1.230517609 | 13 |
| 1.486852078 | 0.974694135 | 1.296829781 | 16 |
| 1.271303757 | 0.883039758 | 0.779884995 | 8 |
| 1.179874056 | 0.9124618 | 0.899264396 | 23 |
| 1.721292922 | 0.974694135 | 0.908446264 | 3 |
| 1.50708932 | 0.974694135 | 0.790506487 | 4 |
| 1.07620358 | 0.9124618 | 0.871911573 | 60 |
| 1.199669427 | 0.974694135 | 0.843180389 | 22 |
| 1.229020768 | 0.974694135 | 0.827179344 | 17 |
| 1.44230072 | 0.974694135 | 0.750335804 | 3 |
| 1.524176419 | 0.974694135 | 0.805277119 | 4 |
| 1.495893842 | 0.974694135 | 0.783073961 | 4 |
| 1.362841175 | 0.974694135 | 0.70965153 | 3 |
| 1.199326659 | 0.974694135 | 0.786035083 | 19 |
| 1.218042591 | 0.974694135 | 0.707099405 | 11 |
| 1.108310722 | 0.974694135 | 0.63600268 | 20 |
| 1.308149609 | 0.974694135 | 0.731413132 | 6 |
| 1.191792129 | 0.974694135 | 0.65427655 | 6 |
| 1.1078681 | 0.694648631 | 0.896456241 | 44 |
| 1.224625474 | 0.974694135 | 0.577818379 | 4 |
| 1.136983932 | 0.974694135 | 0.553888858 | 12 |
| 1.097392165 | 0.974694135 | 0.48769109 | 4 |
| 1.048741592 | 0.974694135 | 0.46033466 | 3 |
| 1.055730993 | 0.974694135 | 0.457003551 | 4 |
| 1.056725596 | 0.974694135 | 0.419995202 | 18 |
| 1.05120283 | 0.974694135 | 0.436324133 | 19 |
| 1.013680701 | 0.974694135 | 0.376599336 | 25 |
| 1.027245496 | 0.974694135 | 0.402489848 | 26 |
| 1.040266061 | 0.974694135 | 0.427606934 | 5 |
| 1.037918959 | 0.974694135 | 0.437828847 | 18 |
| 0.983495428 | 0.931107291 | 0.376659823 | 4 |
| 0.974441283 | 0.974694135 | 0.337970916 | 8 |
| 0.9261052 | 0.788226025 | 0.344539523 | 10 |
| 1.048266132 | 0.974694135 | 0.438413587 | 9 |
| 0.926867494 | 0.974694135 | 0.34983534 | 4 |
| 0.91863718 | 0.943577968 | 0.291922838 | 7 |
| 1.053521682 | 0.850229466 | 0.436394964 | 13 |
| 0.91763082 | 0.943577968 | 0.290294466 | 7 |
| 1.064289192 | 0.694648631 | 0.528708289 | 3 |
| 1.149896005 | 0.756880965 | 0.536929951 | 9 |
| 1.066636324 | 0.694648631 | 0.460406396 | 18 |
| 0.987915836 | 0.850229466 | 0.338715247 | 8 |

| Median Final score | Median Kinase Change | Median Kinase Statistic | SD Kinase Statitistic |
|---|---|---|---|
| 3.673664139 | 0.226304279 | 2.949047483 | NA |
| 3.673664139 | 0.375789851 | 2.912335109 | 0.203763436 |
| 2.935103512 | 0.299158222 | 2.624403059 | 0 |
| 2.594482893 | 0.325927911 | 1.965797985 | 0 |
| 2.594482893 | 0.348389934 | 2.415259119 | 0 |
| 2.527536103 | 0.270422884 | 1.404272006 | 0.195500943 |
| 2.49757288 | 0.245937412 | 1.279045885 | 0.081288801 |
| 2.443215218 | 0.211212879 | 1.864052792 | 0.162568733 |
| 2.258690791 | 0.292021707 | 1.427273142 | 0.157849637 |
| 2.058439678 | 0.299392405 | 1.303648659 | 0.433783417 |
| 1.992422902 | 0.208990028 | 1.465084958 | 0.278299447 |
| 1.957660795 | 0.263154738 | 1.209292637 | 0.131982872 |
| 1.892042289 | 0.23492923 | 1.731885582 | 0.025946613 |
| 1.874378307 | 0.231719882 | 1.683988617 | 0.32755385 |
| 1.860750782 | 0.212170872 | 1.076608856 | 0.027407606 |
| 1.810341279 | 0.26439416 | 1.204907341 | 0.026570064 |
| 1.804432419 | 0.23067854 | 1.225246746 | 0.028450958 |
| 1.76517912 | 0.344395778 | 1.525500259 | 0.144105828 |
| 1.76517912 | 0.270659498 | 1.51240199 | 0.09899656 |
| 1.724274132 | 0.206286107 | 1.422002688 | 0.146623212 |
| 1.684345665 | 0.332619022 | 1.362841175 | 0 |
| 1.660826914 | 0.235891787 | 1.18624875 | 0.09006519 |
| 1.65247484 | 0.285942647 | 1.199538808 | 0.092781072 |
| 1.640240384 | 0.250388275 | 1.126881663 | 0.098354633 |
| 1.638239012 | 0.289942508 | 1.319912178 | 0.068774465 |
| 1.633621597 | 0.279330954 | 1.199849547 | 0.225806892 |
| 1.601226945 | 0.246473055 | 1.136622214 | 0.045047429 |
| 1.546559341 | 0.29604486 | 1.219658632 | 0.014900528 |
| 1.524445026 | 0.246148917 | 1.136983932 | 0 |
| 1.458820291 | 0.251159708 | 1.097392165 | 0 |
| 1.443215218 | 0.139307436 | 1.048741592 | 0 |
| 1.433229677 | 0.142640351 | 1.055730993 | 0 |
| 1.415985564 | 0.24887857 | 1.054587309 | 0.058708356 |
| 1.406492411 | 0.210267696 | 1.054902198 | 0.014546835 |
| 1.383629528 | 0.213735735 | 1.038604116 | 0.045051927 |
| 1.383629528 | 0.213735735 | 1.038604116 | 0.02683873 |
| 1.381408068 | 0.224071818 | 1.036109773 | 0.011755756 |
| 1.374811063 | 0.217983197 | 1.033474434 | 0.093741685 |
| 1.359796919 | 0.252775508 | 1.015977689 | 0.051158121 |
| 1.311936303 | 0.204913065 | 0.98442979 | 0.014982761 |
| 1.311936303 | 0.23106554 | 0.97586273 | 0.23254661 |
| 1.306308218 | 0.202185545 | 0.970278149 | 0.154752606 |
| 1.303530177 | 0.191862098 | 0.914113715 | 0.048312895 |
| 1.288057865 | 0.198890267 | 0.937280162 | 0.055928947 |
| 1.277464792 | 0.16134442 | 1.023420232 | 0.088949381 |
| 1.27572413 | 0.228153143 | 0.927225948 | 0.028785384 |
| 1.223356919 | 0.24974135 | 1.064289192 | NA |
| 1.220432366 | 0.193439256 | 1.144560728 | 0.040899925 |
| 1.206097914 | 0.207181774 | 1.098412867 | 0.060631204 |
| 1.20090769 | 0.283931207 | 0.911833105 | 0.111303479 |
