## additional file 6 for "Infection and herbicide exposure implicate c-Abl kinase in α-Synuclein Ser129 phosphorylation"

**Additional file 6.** The table represents upstream kinases that show induced activities upon rotenone + ponatinib compared to rotenone treatment. Upstream kinase analysis (UKA) was performed on the raw data obtained from Pamgene Serine-Threonine kinase screening. The preliminary analysis has been performed by UKA algorithm. The row data to draw the kinome tree was further filtered using the threshold cut-off for the median final score (kinase score) > 1.2. The kinase statistic represents the log fold change scaled by the noise. The median kinase statistic<0 is considered as inhibited of activity and median kinase statistic>0 is considered as induced activity.

| Kinase Name | Kinase Uniprot ID | Kinase Group | Kinase Family | Max Final score |
|---|---|---|---|---|
| PKN1 | Q16512 | AGC | PKN | 3.673664139 |
| CDK5 | Q00535 | CMGC | CDK | 3.673664139 |
| RPS6KC1 | Q96S38 | OTHER | RSKL | 3.673664139 |
| CDK4 | P11802 | CMGC | CDK | 3.071604148 |
| MAPK7 | Q13164 | CMGC | MAPK | 3.673664139 |
| MAPK11 | Q15759 | CMGC | MAPK | 2.673664139 |
| CDK2 | P24941 | CMGC | CDK | 3.196542884 |
| CDK9 | P50750 | CMGC | CDK | 2.559720787 |
| DAPK3 | O43293 | CAMK | DAPK | 2.448550002 |
| MAPK3 | P27361 | CMGC | MAPK | 3.393618635 |
| CDK3 | Q00526 | CMGC | CDK | 2.559720787 |
| CDK19 | Q9BWU1 | CMGC | CDK | 2.527536103 |
| DCLK3 | Q9C098 | CAMK | DCAMKL | 2.443215218 |
| GSK3A | P49840 | CMGC | GSK | 2.673664139 |
| MAPK10 | P53779 | CMGC | MAPK | 2.393618635 |
| MAPK8 | P45983 | CMGC | MAPK | 2.393618635 |
| PDPK1 | O15530 | AGC | PKB | 2.163169714 |
| CDK1 | P06493 | CMGC | CDK | 2.490528648 |
| NUAK1 | O60285 | CAMK | CAMKL | 2.106127321 |
| MAPK1 | P28482 | CMGC | MAPK | 2.314437389 |
| MAPK9 | P45984 | CMGC | MAPK | 2.217527376 |
| TBK1 | Q9UHD2 | OTHER | IKK | 2.242300375 |
| MAPK14 | Q16539 | CMGC | MAPK | 2.155150199 |
| MAPK13 | O15264 | CMGC | MAPK | 2.092588639 |
| COQ8A | Q8NI60 | Atypical | ABC1 | 2.106127321 |
| ROCK1 | Q13464 | AGC | DMPK | 1.971428747 |
| RPS6KA2 | Q15349 | AGC-CAMK | RSK | 1.978645287 |
| MAPK12 | P53778 | CMGC | MAPK | 2.828566099 |
| DYRK1A | Q13627 | CMGC | DYRK | 1.629006066 |
| CDK6 | Q00534 | CMGC | CDK | 1.813835038 |
| CDKL5 | O76039 | CMGC | CDKL | 2.443215218 |
| CDKL3 | Q8IVW4 | CMGC | CDKL | 1.503402424 |
| DAPK2 | Q9UIK4 | CAMK | DAPK | 1.630190641 |
| CDK13 | Q14004 | CMGC | CDK | 1.682438063 |
| AURKA | O14965 | CAMK | AUR | 1.512805043 |
| RPS6KA6 | Q9UK32 | AGC-CAMK | RSK | 3.071604148 |
| GSK3B | P49841 | CMGC | GSK | 1.501524032 |
| ATR | Q13535 | Atypical | PIKK | 1.524386915 |
| MAP3K14 | Q99558 | STE | STE-Unique | 1.501524032 |
| MYLK3 | Q32MK0 | CAMK | MLCK | 1.828566099 |
| TEX14 | Q8IWB6 | OTHER | NKF5 | 2.527536103 |
| AURKB | Q96GD4 | CAMK | AUR | 1.849550591 |
| RPS6KA1 | Q15418 | AGC-CAMK | RSK | 2.163169714 |
| CHEK1 | O14757 | CAMK | CAMKL | 1.6082888 |
| CILK1 | Q9UPZ9 | CMGC | RCK | 1.35222595 |

| Mean Kinase Statistic | Mean Significance Score | Mean Specificity Score |
|---|---|---|
| 1.502363633 | 0.974694135 | 2.665522227 |
| 1.108351918 | 0.974694135 | 2.531731118 |
| 1.6508964 | 0.974694135 | 2.241030946 |
| 1.144637388 | 0.943577968 | 1.94622968 |
| 0.911474697 | 0.891793987 | 1.777685488 |
| 1.091807797 | 0.974694135 | 1.586333409 |
| 0.902025327 | 0.881345633 | 1.776624267 |
| 1.02554101 | 0.974694135 | 1.459418084 |
| 1.437933541 | 0.749579998 | 1.698970004 |
| 0.858870865 | 0.725764798 | 1.849919652 |
| 0.949078133 | 0.756880965 | 1.560652265 |
| 1.132067877 | 0.974694135 | 1.430978257 |
| 1.26901696 | 0.974694135 | 1.306544615 |
| 1.095189525 | 0.974694135 | 1.276798244 |
| 0.778333146 | 0.694648631 | 1.378321237 |
| 0.786755888 | 0.694648631 | 1.446492327 |
| 1.317612633 | 0.694648631 | 1.402316178 |
| 0.816056687 | 0.694648631 | 1.383454014 |
| 1.172686187 | 0.749579998 | 1.312225819 |
| 0.836159278 | 0.694648631 | 1.330181291 |
| 0.8016902 | 0.694648631 | 1.28415477 |
| 1.013880304 | 0.819113299 | 1.046557656 |
| 0.789539341 | 0.850229466 | 0.914402928 |
| 0.795684828 | 0.694648631 | 1.137490847 |
| 0.920613423 | 0.583063944 | 1.111450646 |
| 1.01572211 | 0.749579998 | 0.958605327 |
| 0.673326309 | 0.608150577 | 0.789665103 |
| 0.868733305 | 0.787997132 | 1.218664743 |
| 0.985972027 | 0.749579998 | 0.876160635 |
| 0.879896029 | 0.694648631 | 0.937346319 |
| 0.919307498 | 0.716916199 | 0.985269846 |
| 0.790801306 | 0.974694135 | 0.528708289 |
| 0.813999778 | 0.673980817 | 0.716200371 |
| 0.827118681 | 0.834671383 | 0.645010022 |
| 0.86750556 | 0.694648631 | 0.779988211 |
| 0.796360362 | 0.677187564 | 1.01858287 |
| 0.803444255 | 0.679937609 | 0.752066039 |
| 0.801083813 | 0.694648631 | 0.7408822 |
| 0.866760553 | 0.694648631 | 0.733734252 |
| 0.866590424 | 0.741322881 | 0.716910548 |
| 1.010074013 | 0.806666832 | 0.997982789 |
| 0.961292605 | 0.59167148 | 0.863811567 |
| 0.708934702 | 0.606382502 | 0.771159883 |
| 0.754102054 | 0.694648631 | 0.66129894 |
| 0.804250181 | 0.694648631 | 0.573728077 |

| Mean peptide set size | Median Final score | Median Kinase Change |
|---|---|---|
| 7 | 3.673664139 | 0.2079763 |
| 12 | 3.372634143 | 0.221392087 |
| 3 | 3.134073516 | 0.283430455 |
| 7 | 2.974694135 | 0.220991622 |
| 13 | 2.828566099 | 0.229426096 |
| 6 | 2.559720787 | 0.258724082 |
| 17 | 2.548520595 | 0.182230749 |
| 8 | 2.469544156 | 0.210492207 |
| 3 | 2.448550002 | 0.213757685 |
| 20 | 2.439376125 | 0.211423451 |
| 11 | 2.393618635 | 0.195180307 |
| 5 | 2.372634143 | 0.213446884 |
| 3 | 2.242598449 | 0.204617803 |
| 5 | 2.196784293 | 0.195022133 |
| 25 | 2.163169714 | 0.186784166 |
| 26 | 2.163169714 | 0.186784166 |
| 3 | 2.13834613 | 0.195598806 |
| 20 | 2.092588639 | 0.189300759 |
| 4 | 2.069851988 | 0.208895231 |
| 18 | 2.013407393 | 0.192447315 |
| 19 | 1.995678626 | 0.184552823 |
| 5 | 1.91649738 | 0.208859707 |
| 14 | 1.854120204 | 0.188317919 |
| 18 | 1.802554028 | 0.186091363 |
| 8 | 1.784098187 | 0.202407314 |
| 4 | 1.700361975 | 0.18732604 |
| 23 | 1.637743779 | 0.171585505 |
| 11 | 1.630190641 | 0.217618613 |
| 3 | 1.625740633 | 0.234037282 |
| 7 | 1.6082888 | 0.177704327 |
| 6 | 1.5703912 | 0.185138718 |
| 3 | 1.503402424 | 0.365754205 |
| 5 | 1.479804783 | 0.248870913 |
| 4 | 1.474939445 | 0.195873763 |
| 5 | 1.474540543 | 0.170802313 |
| 16 | 1.460447164 | 0.217392437 |
| 7 | 1.439376125 | 0.172279662 |
| 7 | 1.425135686 | 0.187048891 |
| 4 | 1.41589503 | 0.114522061 |
| 4 | 1.393640351 | 0.189439851 |
| 4 | 1.376585296 | 0.162494635 |
| 4 | 1.374728716 | 0.171388373 |
| 22 | 1.331800516 | 0.193293162 |
| 9 | 1.325432773 | 0.152350767 |
| 3 | 1.321736628 | 0.215384522 |
